# Native and invading yellow starthistle (*Centaurea solstitialis*) microbiomes differ in composition and diversity of bacteria

**DOI:** 10.1101/119917

**Authors:** Patricia Lu-Irving, Julia Harenčár, Hailey Sounart, Shana R Welles, Sarah M Swope, David A Baltrus, Katrina M Dlugosch

## Abstract

- Invasive species could benefit from introduction to locations with favorable species interactions. Microbiomes are an important source of interactions that vary across regions. We examine whether bacterial communities could explain more favorable microbial interactions in highly invasive populations of yellow starthistle.
- We sequenced amplicons of prokaryotic 16S rRNA genes to characterize bacterial community composition in the phyllosphere, ectorhizosphere, and endorhizosphere of plants from seven invading populations in California, USA and eight native populations in Europe. We tested for differentiation of microbiomes by geography, plant compartment, and plant genotype.
- Bacterial communities differed significantly between native and invaded ranges within plant compartments, with consistently lower diversity in plants from the invaded range. Genera containing known plant pathogens also showed lower diversity in invaded range plants. The diversity of bacteria in roots was positively correlated with plant genotype diversity within both ranges, but this relationship did not explain microbial differences between ranges.
- Our findings reveal changes in the composition and diversity of bacterial interactions in invading plants, consistent with observations of altered soil interactions in this invasion. These results call for further study of the sources of variation in microbiomes and the potential for bacteria to facilitate invasion success.

## INTRODUCTION

Humans continue to transport plant species around the globe, and increasing numbers of these translocations result in the invasive expansion of non-native species into recipient communities (Lonsdale, 1999; Butchart *et al.*, 2010; Essl *et al.*, 2011; Ellis *et al.*, 2012). While there are undoubtedly many reasons that species introductions lead to invasions, there is growing evidence that novel species interactions may facilitate the invasive spread of populations (Mitchell *et al.*, 2006; Pearson *et al.*, 2018). Initially, hypotheses about the contribution of species interactions to invasions focused on the potential for non-native species to escape from above-ground herbivores, which are easily observed (Keane & Crawley, 2002), though it is not clear that herbivore escape is a frequent mechanism of invasion (Maron & Vila, 2001; Levine *et al.*, 2004; Agrawal *et al.*, 2005; Felker-Quinn *et al.*, 2013). More recently, there has been increasing recognition that microbial taxa above- and below-ground can have large effects on plant fitness, both positive and negative, and could thus determine whether invasive plants benefit from novel species interactions (Callaway *et al.*, 2004; Colautti *et al.*, 2004; Torchin & Mitchell, 2004; Agrawal *et al.*, 2005; Mitchell *et al.*, 2006; Kulmatiski *et al.*, 2008; van der Putten *et al.*, 2016; Faillace *et al.*, 2017). Plant-associated microbial communities have been historically difficult to observe, however, and studies that leverage newly-available tools to quantify differences in these interactions across native and invading populations are needed to test hypotheses for invasion success (Dawson & Schrama, 2016).

Microbial communities have emerged as particularly likely candidates for facilitating invasions. Although many interactions between plants and microbes can be beneficial, soil microbial communities often appear to have negative net effects on plant fitness which may become more negative over time, e.g., via plant-soil feedbacks (Bever, 2003; Reinhart & Callaway, 2006; Kulmatiski *et al.*, 2008; Petermann *et al.*, 2008). These interactions between plants and their microbiomes can vary over space and environment (Nemergut *et al.*, 2013; van der Putten *et al.*, 2013; terHorst & Zee, 2016; Lankau & Keymer, 2018), creating opportunities for introduced plants to escape negative interactions that might characterize their native ranges. Moreover, reductions in microbial diversity are occurring in response to environmental change and human disturbances, and lowered microbial diversity may reduce the resistance of ecosystems to invasion (Schnitzer *et al.*, 2011; Wagg *et al.*, 2014; Dawson & Schrama, 2016; van der Putten *et al.*, 2016).

Invasive plant species have provided some of the best evidence to date that microbial interactions can be locally evolved, and can vary considerably over geographic regions (Rout & Callaway, 2012). Introduced plants have been shown to vary in their response to soil communities from their native and invaded ranges, and there are now many examples of more favorable interactions between plants and soil from their invaded range, consistent with escape from enemies or gain of mutualists during invasion (Reinhart *et al.*, 2003; Callaway *et al.*, 2004; Mitchell *et al.*, 2006; Engelkes *et al.*, 2008; Kulmatiski *et al.*, 2008; Maron *et al.*, 2014; van der Putten *et al.*, 2016). Plant-microbe interactions which provide relative benefits to invasive species can be explained by reduced negative effects of key microbial pathogens, increased direct beneficial effects of mutualistic taxa, or increased indirect benefits from taxa that affect competitors more negatively than they do the invader (Dawson & Schrama, 2016). It is also possible that invaders could benefit from a reduced diversity of enemy interactions, as a result of an associated reduction in ecological costs that derive from simultaneously deploying different defense responses against many different enemies (Agrawal, 2007; Colautti & Lau, 2015; Smakowska *et al.*, 2016). These hypotheses all require that there are differences in the microbial communities associated with invading vs. native plants, specifically changes in taxonomic composition, and the loss or gain of groups with pathogenic or mutualistic effects (Dawson & Schrama, 2016).

Here we conduct one of the first comparisons of plant microbiomes between invading populations and populations in their native source region. We survey plant-associated microbial communities in the highly invasive forb yellow starthistle (*Centaurea solstitialis*). Yellow starthistle is native to a wide region of Eurasia and was introduced from western Europe to South America in the 1600’s and North America in the 1800’s as a contaminant of alfalfa seed (Gerlach, 1997; Barker *et al.*, 2017). This herbaceous annual is a colonizer of grassland ecosystems, and is often cited as one of the ‘10 Worst Weeds of the West’ in North America (DiTomaso & Healy, 2007). Its extensive invasion of California in the USA (>14 million acres; Pitcairn *et al.*, 2006) is particularly well-studied, and invading genotypes in this region have evolved to grow larger and produce more flowers than plants in the native range, suggesting a shift in resource allocation that has favored invasiveness (Widmer *et al.*, 2007; Eriksen *et al.*, 2012; Dlugosch *et al.*, 2015). Previous research has demonstrated that yellow starthistle throughout all of its native and invaded ranges experiences net fitness reductions when grown with its local soil communities (Andonian *et al.*, 2011, 2012; Andonian & Hierro, 2011). These studies have also indicated that this negative interaction is weaker (more favorable) in California, raising the possibility that changes in the microbial community have promoted an aggressive invasion.

We sample microbial communities associated with leaves (phyllosphere and endosphere) and roots (ectorhizosphere and endorhizosphere) of yellow starthistle plants in both the California invasion and native regions in Europe. Previous experiments with fungicide treatments have shown that plant-soil interactions between yellow starthistle and fungi in California are *more* negative (less favorable) than those in the native range, inconsistent with a role for fungi in beneficial species interactions in this invasion (Hierro *et al.*, 2017). Here, we focus on bacterial communities, using high-throughput sequencing of prokaryotic ribosomal 16S amplicon sequences to quantify taxonomic composition and diversity of bacteria in yellow starthistle microbiomes. Microbial communities are known to differ among plant compartments (Bulgarelli *et al.*, 2013) and to be influenced by individual plant genotype (Lundberg *et al.*, 2012; Peiffer *et al.*, 2013; Bodenhausen *et al.*, 2014), and so we explicitly test for differences in the microbiomes of native and invaded range plants relative to the influence of both plant compartment and plant genotype.

## MATERIALS AND METHODS

### Study species

Yellow starthistle (*Centaurea solstitialis* L., Asteraceae) is an obligately outcrossing annual plant, diploid throughout its range (Irmina *et al.*, 2017). Plants form a taproot and grow as a rosette through mild winter and/or spring conditions, bolting and producing flowering heads (capitula) throughout the summer. The species is native to Eurasia, where distinct genetic subpopulations have been identified in Mediterranean western Europe, central-eastern Europe, Asia (including the Middle East), and the Balkan-Apennine peninsulas (Barker *et al.*, 2017). The invasion of California as well as invasions in South America appear to be derived almost entirely from western European genotypes (Fig. 1; Barker *et al.*, 2017).

**Fig.1.**
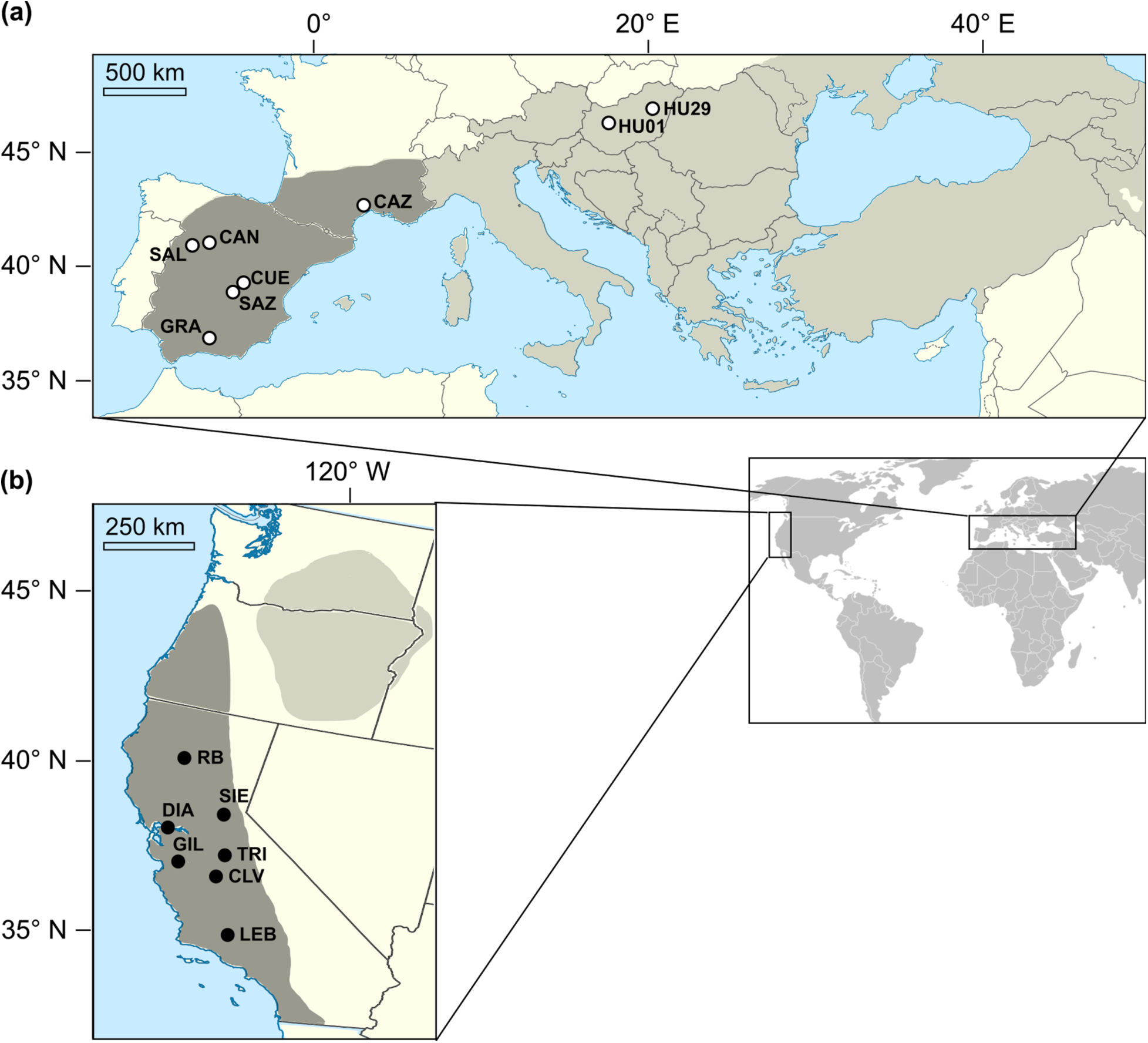
The distribution (gray) of yellow starthistle and sampling sites (circles) for this study. Maps detail the native range in Eurasia (a) and the invasion of western North America (b). Previous work has indicated that western Europe is the source for the severe invasion of California, USA (both in dark shading; Barker et al. 2017). Sampling included seven locations in California (b, filled circles), six locations in western Europe and an additional two locations in eastern Europe (a, open circles).

### Sample collection

Fifteen populations of yellow starthistle were sampled for microbial communities: seven sampling sites across the invasion of California, six in western Europe, and two in eastern Europe (Fig. 1; Supporting Information Table S1). At each location, plants were sampled every meter (or to the nearest meter mark) along a 25 meter transect, to yield microbial samples from 25 individual plants per population. Individuals in rosette or early bolting stages were preferentially selected. In one population (HU29), low plant density yielded 20 individuals along the 25 meter transect. Using sterile technique, plants were manually pulled and each individual sampled using modified versions of protocols by Lundberg *et al.* (2013) and Lebeis *et al.* (2015) as described below. Plants were pressed and dried after sampling, and submitted to the University of Arizona Herbarium (ARIZ; Supporting Information Table S1).

#### Phyllosphere and ectorhizosphere

One to three basal, non-senescent leaves were collected from each plant, as well as the upper 2-5 cm of the taproot, together with accompanying lateral roots (excess soil was brushed or shaken off). Leaf and root samples were placed in individual 50 ml tubes containing 25 ml of sterile wash solution (45.9 mM NaH_2_PO_4_, 61.6 mM Na_2_HPO_4_, 0.1% Tween 20). Tubes were shaken by hand for one minute (timed). Leaf and root samples were then removed and stored on ice in separate tubes (leaves in empty tubes, roots in tubes containing 10 ml of wash solution) until further processing. Wash samples were stored on ice during transport, then refrigerated at 4°C. Phyllosphere and ectorhizosphere washes were pooled across all (20 or 25) plants at a location, then centrifuged at 2,200 g at 4°C for 15 minutes. Supernatants were discarded, and pellets were air-dried and stored at −20°C until DNA extraction.

#### Leaf endosphere

Leaves were surface sterilized by submerging in bleach solution (10% commercial bleach, 0.1% tween 20) for two minutes. Leaves were then rinsed in distilled water, patted dry using a kimwipe, and sealed in individual sterile surgical envelopes (Fisherbrand #01- 812-50). Envelopes were kept in silica gel desiccant until leaf tissue was completely dry, then stored at room temperature until DNA extraction.

#### Endorhizosphere

Roots were further washed by shaking in 10 ml of wash solution until visible residual soil was removed. Washed roots were stored and dried as described above for leaves.

#### Controls

At each collection site, a tube of sterile wash solution was left uncapped while sampling plants. Disinfected tools were periodically agitated in the blank wash tube before sterilization and use for the next sample collection. For each population, rinse water and wipes used to process tissue samples were represented in controls by rinsing and wiping flame-sterilized forceps, then agitating the forceps in the blank wash tube. Controls were stored and processed in the same manner as phyllosphere and ectorhizosphere samples.

### DNA extraction

Extractions were carried out using sterile technique in a laminar flow hood. Leaf and root DNA were extracted from tissue pooled by sampling site (15 total populations), and as individual plant samples from 8 plants from each of 10 populations (80 total individuals). For pooled tissue extractions, equal sections of leaf tissue (50 mm^2^) and root tissue (12.5 mm^3^ plus 10 mm of lateral roots) were collected from each individual sample per location and pooled prior to extraction. Control (blank) samples were collected for each batch of extractions by swabbing tools and surfaces, then extracting DNA from the swab head.

All DNA samples were extracted using the MO BIO PowerSoil kit (MO BIO Laboratories, Inc.). Phyllosphere and ectorhizosphere DNA was extracted from up to 0.25 g of wash pellets following the standard kit protocol. Leaf and root tissues were ground to powder or sawdust consistency in liquid nitrogen using sterile mortars and pestles. Leaf and root DNA was extracted from 20 mg (leaf) or 100 mg (root) of ground tissue with the following modification to the standard protocol: tissue was incubated at 65°C for 10 minutes in extraction buffer, then vortexed for 1 minute, followed by a second 10 minute incubation (as described under “alternative lysis methods” in the kit protocol). Control DNA was extracted by placing whole swab heads directly into extraction tubes. Extracted DNA was eluted in PCR-grade water and stored at −20°C pending library preparation.

### Library preparation and sequencing

To remove secondary compounds inhibiting PCR, DNA extracted from root and leaf tissue (together with corresponding blanks) was purified using a ZR-96 genomic DNA clean-up kit (Zymo Research). All DNA concentrations were quantified using a Qubit fluorometer high-sensitivity assay for double-stranded DNA (Invitrogen), and standardized to equimolar amounts.

Library preparation followed a dual-index approach (Kozich *et al.*, 2013) using a two-step PCR protocol as follows. In the first step (target-specific PCR), the V4 region of the 16S rRNA gene was amplified using target specific primers (515F and 806R; Caporaso *et al.*, 2011) appended with common sequence (CS) tags through a linker sequence which varied from two to five nucleotides in length. Target-specific PCR was carried out using Phusion Flash master mix (Thermo Scientific) in 25 µl reaction volume in a Mastercycler pro thermocycler (Eppendorf) under the following conditions: 25 cycles of 1 s at 98°C, 5 s at 78°C, 5 s at 57°C, 15 s at 72°C. Products were visualized on an agarose gel and diluted by up to 1:15 (depending on yield); 1 µl of diluted product was then used as template in the second step (barcode-adapter attachment PCR). Using reagents and equipment as described above, barcoded primer pairs incorporating Illumina P5 and P7 adapters were used to amplify products from target-specific PCR in 25 µl reaction volumes under the following conditions: 10 cycles of 1 s at 98°C, 5 s at 78°C, 5 s at 51°C, 15 s at 72°C. Barcoded amplicons were quantified by fluorometry, pooled in equimolar amounts, cleaned, and submitted to the University of Idaho’s IBEST Genomic Resources Core for QC and sequencing. Amplicons were multiplexed to use half the capacity of one 2 × 300 bp run on an Illumina MiSeq platform. Raw sequence data are deposited in the NCBI Short Read Archive under accession number XXXXXX [pending submission].

Peptide nucleic acid clamps (PNAs) were included in both PCR steps of library preparation to block amplification of plant chloroplast and mitochondrial 16S as recommended by Lundberg *et al.* (2013). Clamp sequences published by Lundberg *et al.* (2013) were compared with chloroplast and mitochondrial 16S sequences from yellow starthistle and three other species of Asteraceae with published organellar genomes (*Centaurea diffusa, Helianthus annuus, Lactuca sativa*). We found a single nucleotide mismatch between the Asteraceae chloroplast 16S and the plastid PNA sequences, and designed an alternative plastid PNA specific to the Asteraceae sequence (5’—GGCTCAACTCTGGACAG—3’) (Fitzpatrick *et al.*, in revision). All samples for this study were amplified using the plastid PNA of our design, together with the mitochondrial PNA published by Lundberg *et al.* (2013). To gauge the effectiveness of our alternative PNA, two duplicate samples were processed using both PNAs published by Lundberg *et al.* (2013).

### Identification of operational taxa and potential plant pathogens

Demultiplexed paired reads were merged and quality filtered using tools from the USEARCH package version 9.0.2132 (Edgar & Flyvbjerg, 2015). Merged reads were truncated to uniform length and primer sequences were removed using a combination of the seqtk toolkit version 1.2 (github.com/lh3/seqtk) and a custom script. The UPARSE pipeline (Edgar, 2013) implemented in the USEARCH package was used for further data processing and analysis: unique sequences were identified, and those represented only once or twice in the processed read set were discarded as likely PCR or sequencing errors. Remaining sequences were clustered into operational taxonomic units (OTUs) at a 97% threshold, chimeras were filtered out, and per-sample OTU read counts were tabulated using the UPARSE-OTU algorithm. Assignment of OTUs to nearest taxonomic match in the Greengenes database (McDonald *et al.*, 2012) was carried out using the UCLUST algorithm implemented in QIIME version 1.9.1 (Caporaso *et al.*, 2010; Edgar, 2010). Data were further processed using tools from the QIIME package: reads mapping to chloroplast and mitochondrial OTUs were removed, and samples were rarefied by plant compartment. Rarefaction levels were chosen to reflect the distribution of read counts per sample within plant compartments, subsampling to the minimum number of reads necessary to include all samples except those that were outliers for low read count.

Taxa known to contain plant pathogens were identified using the FAPROTAX database (version 1.1, (Louca *et al.*, 2016). A list of all genera included under the “plant pathogen” functional category was used to filter our OTU tables by taxonomic assignment. While pathogen-containing genera are likely to include both pathogenic and non-pathogenic strains (Newton et al., 2010), our pathogen-containing OTU dataset should be enriched for potential plant pathogens, relative to the full dataset.

### Microbial community analyses

All statistical analyses were performed in R (R Core Team, 2015). We evaluated differences in bacterial community composition between plant compartments, and between native and invaded ranges within plant compartments, by performing non-metric multidimensional scaling (NMDS) using the R packages vegan (Oksanen *et al.*, 2016) and MASS (Venables & Ripley, 2002). Individual plant samples or samples pooled within sampling site provided replicates in these comparisons. Ordinations were based on Bray-Curtis distances, and were performed using a two-dimensional configuration to minimize stress, using Wisconsin double standardized and square root transformed data, with expanded weighted averages of species scores added to the final NMDS solution. Significant differences among plant compartments and between native and invaded samples were assessed using the envfit function in vegan. Ellipses were drawn on NMDS plots using the vegan function ordiellipse, representing 95% confidence limits of the standard error of the weighted average of scores.

We further explored the underlying correlates of bacterial community variation using Principal Components Analysis (PCA; using R function prcomp) for samples from native and invaded ranges within each plant compartment. Prior to performing the PCA, we performed Hellinger’s transformation to minimize the influence of OTUs with low counts or many zeros (Legendre & Legendre, 1998; Legendre & Gallagher, 2001; Ramette, 2007). We then identified the OTUs with the highest loading on the dominant PC axis of variation by examining the matrix of variable loadings produced by prcomp. The OTU composition of samples pooled by sampling site (phyllosphere, ectorhizosphere, and endorhizosphere samples; hereafter ‘pooled samples’) was visualized using a heatmap generated in ggplot2 (Wickham, 2009), and samples were hierarchically clustered by Bray-Curtis dissimilarity (hclust function in R) using McQuitty’s method (McQuitty, 1966).

We compared the diversity of OTUs between the native and invaded range for each plant compartment using richness (R), evenness (Pielou’s J; Pielou, 1966), and their combined effects via the Hill series exponent e^H’^ (Hill, 1973) of the Shannon diversity index (H’; Shannon, 1948). Again, individual plant samples or samples pooled by sampling site provided replicates in these comparisons. Diversity values were calculated using the packages vegan and iNEXT (Hsiegh *et al.*, 2016), and compared between native and invaded ranges using a nonparametric Kruskal-Wallis rank sum test on rarefied read counts. For plant tissue samples that included multiple individuals per site, we compared diversity between regions using a nested ANOVA with fixed effects of region and population nested within region, and among sites using a post hoc Kruskal-Wallis test within regions. We conducted these comparisons on a dataset including all OTUs, and a reduced dataset including only OTUs assigned to genera with known plant pathogens according to the FAPROTAX database, as described above.

Finally, we examined the influence of plant genotype on microbial composition. Our geographic regions correspond to genetically differentiated subpopulations, and within these regions, our study sites are also known to vary in plant genetic diversity (Barker *et al.*, 2017). Microbial diversity estimates (e^H’^) for each plant compartment were predicted using linear models that included fixed effects of plant genetic diversity, region (native vs. invaded), and the interaction between these two effects. Measurements of plant genetic diversity at each of our sampling sites were obtained from previously published genome-wide marker analyses by Barker and colleagues (Barker *et al.*, 2017), calculated as the average proportion of pairwise nucleotide differences between alleles (π) at variable positions across the yellow starthistle genome.

## RESULTS

### Sequencing and data processing

Sequencing yielded 9,672,898 read pairs, of which 6,217,852 remained after merging and quality control; these were 253 bp in length after removing artificial and primer sequences. The number of raw read counts per sample ranged from 16 to 306,200 with a median of 21,964. Analysis of the merged and processed reads resulted in 4,014 OTUs, of which 60 were identified as plastid or mitochondrial, and 428 were unidentifiable (11%). Of the remaining 3,526, 206 were identified to species (6%), 1,084 to genus (27%), and 2,229 to family (56%). A total of 103 OTUs (3%) were identified as members of the 49 genera with known plant pathogens in the FAPROTAX v.1.1 database.

Sequence reads representing yellow starthistle chloroplast and mitochondrial 16S accounted for 40% and 1% of all reads, respectively. Amplification of host chloroplast in samples using the Asteraceae-specific plastid PNA was reduced by up to 51% compared with the Lundberg *et al.* (2013) PNA (Supporting Information Table S2). Despite PNA blocking activity, 83% of the total reads from leaf endosphere samples were yellow starthistle chloroplast sequences. After removal of chloroplast and mitochondrial reads, remaining read counts for most leaf endosphere samples were low relative to controls (Supporting Information Fig. S1), so no further analysis of leaf endosphere bacterial communities was performed.

Rarefaction levels (chosen to reflect the minimum number of reads per sample by compartment, not including outliers) were 18,000 reads per sample for phyllosphere, 17,000 for ectorhizosphere, and 5,000 for endorhizosphere samples (Supporting Information Fig. S1). These levels were also higher than nearly all control samples. Rarefaction cutoffs resulted in the exclusion of five non-control samples which were outliers for low read count: one phyllosphere (DIA), one ectorhizosphere (SAZ), and three individual endorhizosphere samples (two from SAZ; one from SIE). An NMDS ordination of all un-rarefied samples showed that the controls clustered together and were clearly differentiated from all samples in all plant compartments other than the leaf endosphere (Supporting Information Fig. S2).

### Microbial community analyses

Results from NMDS ordinations indicated that bacterial communities differed overall among the phyllosphere, ectorhizosphere, and endorhizosphere compartments (Fig. 2a; stress = 0.14, *P* = 0.001). Within compartments, NMDS further revealed significant differences between native and invaded range endorhizosphere samples (Fig. 2b; stress = 0.16, *P* = 0.001) and ectorhizosphere samples (*P* = 0.001). Native and invaded range phyllosphere samples differed with marginal significance (*P* = 0.05). Clustering analyses within the phyllosphere and ectorhizosphere compartments consistently grouped invaded range samples together, as well as samples from the source region in western Europe (Supporting Information Fig. S3). Native range samples from eastern Europe (HU01 and HU29) clustered together in these compartments but were variable in their relationship to the other regions. Endorhizosphere samples pooled by location showed less consistent clustering by range.

**Fig 2.**
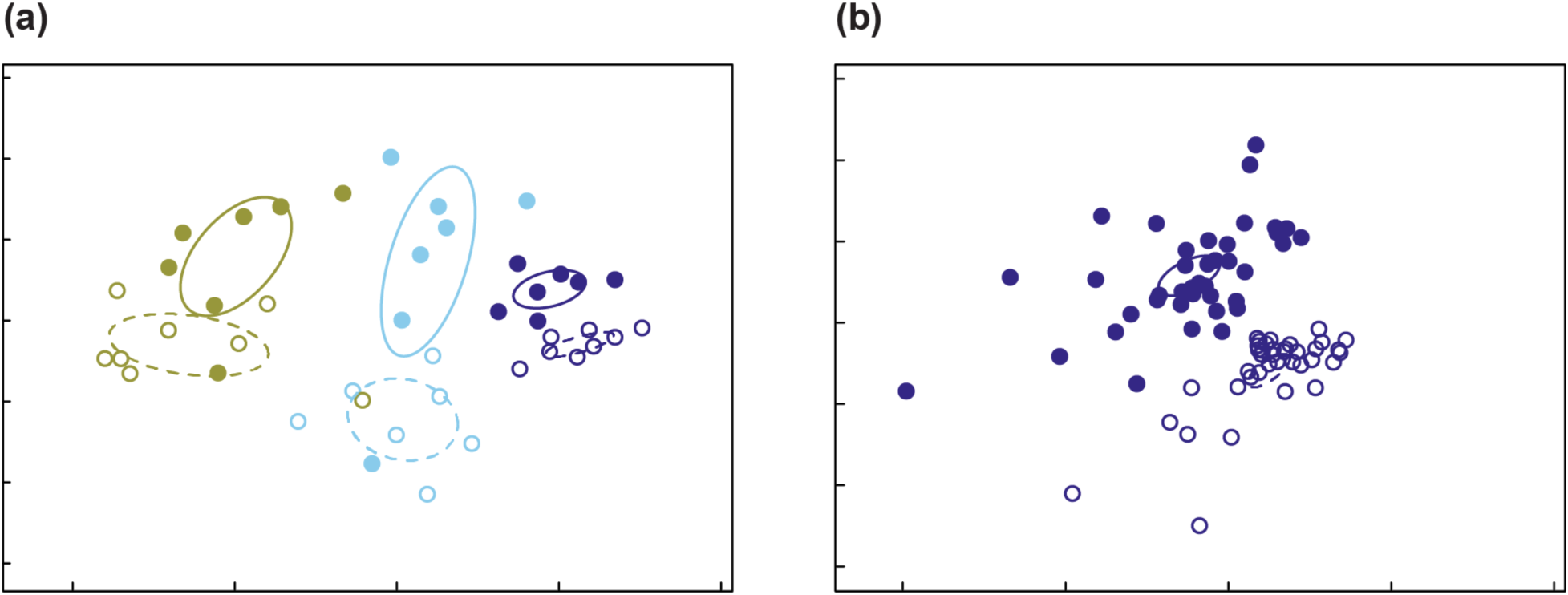
NMDS plots of bacterial OTU composition in phyllosphere (green), endorhizosphere (light blue), and ectorhizosphere (dark blue) samples from native (open symbols) and invaded (closed symbols) ranges. Plotted are a) pooled samples for each sampling location, showing overall separation by range within compartment (stress = 0.14), and b) whole root samples for individual plants within native and invading populations (stress = 0.16). Ellipses indicate 95% confidence intervals for samples grouped by range (native range: dashed lines; invaded range: solid lines).

The dominant phyla among all bacterial communities were Proteobacteria, Actinobacteria, Bacteroidetes, and Firmicutes (Fig. 3). Principal component analyses suggested that the strongest contributions to changes in bacterial community composition between the native and invaded ranges were made by shifts in the representation of *Bacillus* (Firmicutes), *Chryseobacterium* (Bacteroidetes), and the Proteobacteria taxa *Erwinia, Pseudomonas*, and Xanthomonadaceae (Supporting Information Table S3; Fig. S4). All of these taxa other than *Chryseobacterium* include known plant pathogens in the FAPROTAX v.1.1 database (Louca *et al.*, 2016).

**Fig 3.**
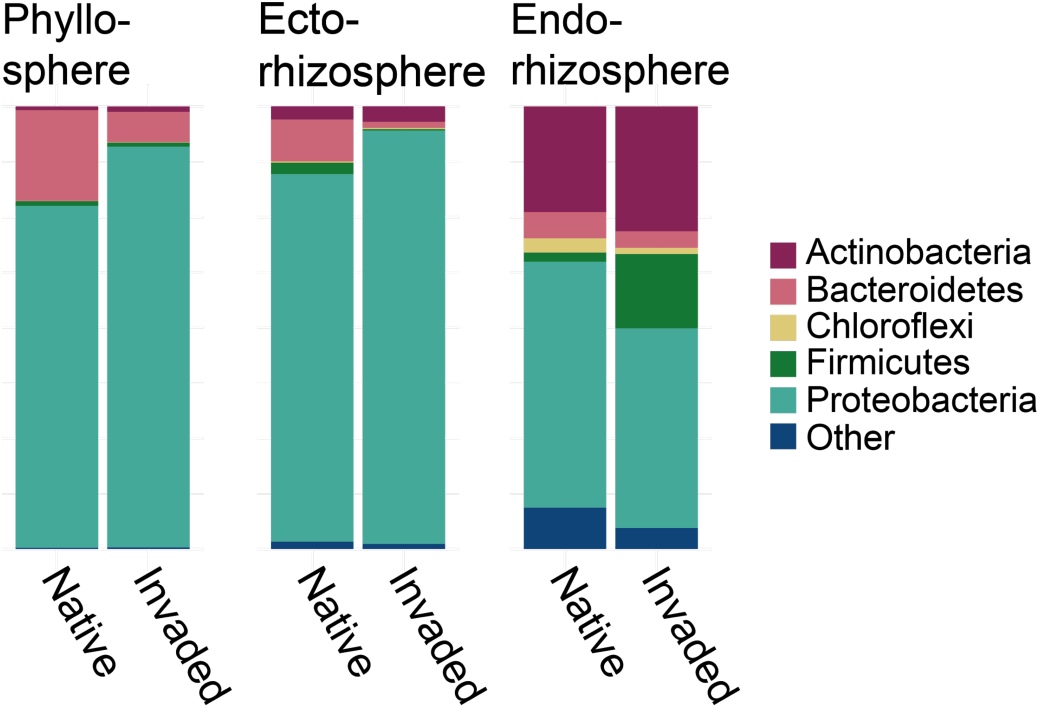
Relative abundance of (proportion of reads mapping to) phyla in yellow starthistle phyllosphere, ectorhizosphere, and endorhizosphere samples from native and invaded ranges.

In general, bacterial OTUs showed a pattern of lower median richness (R), evenness (J), and diversity (e^H’^) for plants from invaded range sites in all compartments, with the exception of richness in the phyllosphere (Figs 4, 5). Within the phyllosphere, invaders were not significantly different in richness (χ^2^_1_ = 1.67, *P* = 0.20), were significantly lower in evenness (χ^2^_1_ = 8.07, *P* = 0.005), and were marginally lower in Hill diversity (χ^2^_1_ = 3.75, *P* = 0.05), relative to native range plants. Similarly, the ectorhizosphere of invaders was not significantly different in richness (χ^2^_1_ = 0.69, *P* = 0.41), and was significantly lower in both evenness (χ^2^_1_ = 5.0, *P* = 0.03) and diversity (χ^2^_1_ = 6.21, *P* = 0.01). In contrast, endorhizosphere samples pooled by site did not differ significantly in evenness (χ^2^_1_ = 2.26, *P* = 0.13), but were marginally significantly lower in richness (χ^2^_1_ = 3.43, *P* = 0.06) and diversity (χ^2^_1_ = 3.01, *P* = 0.08). Nested ANOVA of individual endorhizosphere samples indicated strongly significant reductions in richness, evenness, and diversity in invading plants (Fig 5; fixed effect of region: all *P* < 0.001). For individual endorhizosphere samples, populations did not differ significantly in any metrics within native/invaded regions, with the exception of significantly higher evenness in plants at site SIE relative to site TRI in the invaded range.

**Fig 4.**
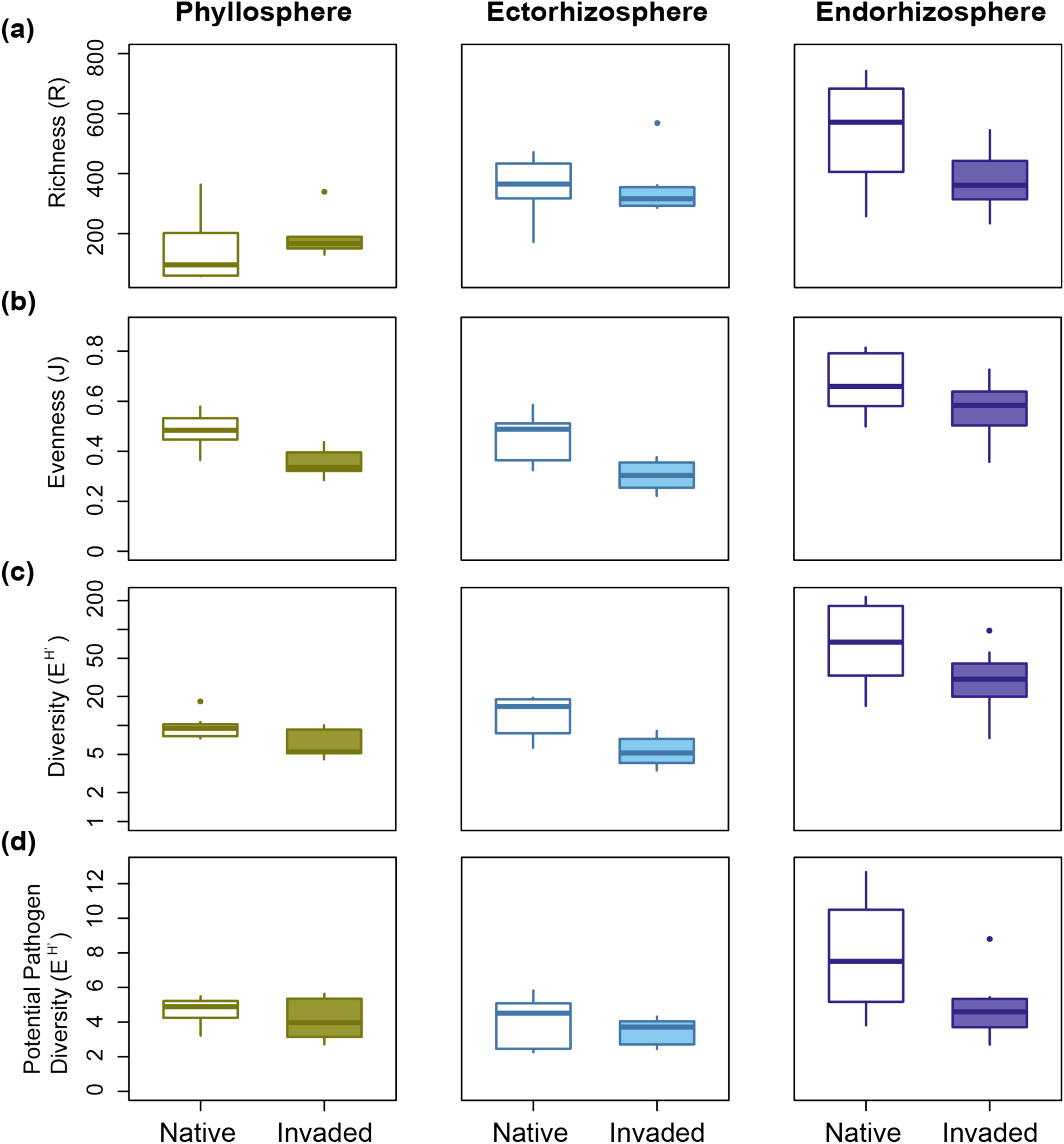
Distributions of OTU (a) richness, (b) evenness, and (c,d) diversity (e^H’^) among samples (pooled plants) from each location in the native and invaded ranges for phyllosphere, ectorhizosphere, and endorhizosphere compartments. (a-c) show values for all OTUs and (d) shows values based on OTUs from known pathogen-containing genera. Significant differences from Kruskal-Wallis tests are indicated with asterisks: \**P* < 0.05, ^(^*^)^*P* < 0.1.

**Fig 5.**
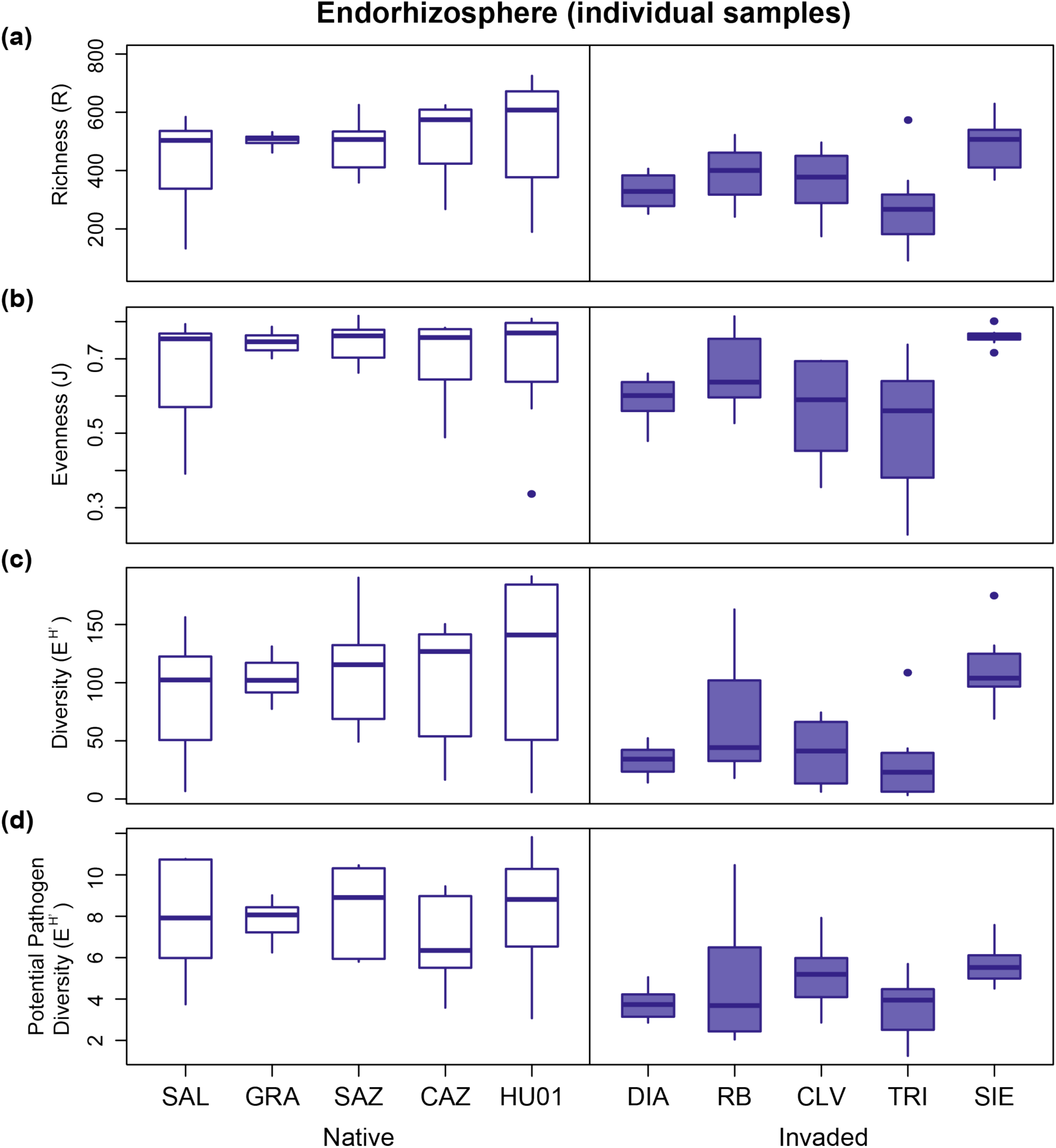
Distributions of endorhizosphere OTU (a) richness, (b) evenness, and (c,d) diversity (e^H’^) among individual plants at each location in the native and invaded ranges. (a-c) show values for all OTUs and (d) shows values based on OTUs from known pathogen-containing genera. Significant differences from Kruskal-Wallis tests are indicated with asterisks: \**P* < 0.05, ^(^*^)^*P* < 0.1.

Filtering the rarefied datasets for genera containing known plant pathogens resulted in 49 phyllosphere OTUs, 69 ectorhizosphere OTUs, and 88 endorhizosphere OTUs enriched for potential plant pathogens. *Pseudomonas* and *Erwinia* were among the most common pathogen-containing genera encountered in each plant compartment in both ranges. The phyllosphere also included a high frequency of *Janthinobacterium*, with a relative increase in *Serratia* in the invaded range. In the ectorhizosphere, *Serratia* was common in both ranges, but invading plants showed a large relative decline in *Erwinia* and increase in *Pseudomonas*. In the endorhizosphere, invading plant harbored less *Pseudomonas* and more *Bacillus* and *Streptomyces* than native plants (Supporting Information Fig. S5). Diversity of these OTUs showed similar trends to total diversity, with lower median values in invaded range root compartments. No differences between regions were statistically significant for the phyllosphere (χ^2^_1_ = 0.60, *P* = 0.44) or ectorhizosphere (χ^2^_1_ = 0.49, *P* = 0.48). For the endorhizosphere samples pooled by site, significantly lower diversity was indicated in the invaded range (χ^2^_1_ = 4.34, *P* = 0.04). For individual endorhizosphere samples, nested ANOVA also indicated significantly lower diversity in the invaded range (*P* < 0.0001), and no significant differences among populations within regions.

The diversity of bacteria in root compartments showed evidence of associations with plant genetic diversity at sampling sites. A linear model predicting microbial diversity (e^H’^) from plant genetic diversity was significant for endorhizosphere samples pooled by site (Fig. 6; *F*_(2,12)_ = 5.89; *P* = 0.02; r^2^_adj_ = 0.41), with significant main effects of both plant genetic diversity (*P* = 0.03) and region (native vs. invaded; *P* = 0.006). The interaction between these two effects was not significant (*P* = 0.71) and was removed from the final model. This same pattern was marginally significant when using only OTUs from pathogen-containing genera in the endorhizosphere samples (effect of plant genetic diversity: *P* = 0.08). Similar linear models did not identify significant effects of plant genetic diversity when predicting the median diversity of individual plant endorhizosphere samples (*P* = 0.74), or diversity in the phyllosphere (*P* = 0.35). There was a marginally significant positive effect of plant genetic diversity on diversity in the ectorhizosphere (*P* = 0.08) again in addition to the effect of region (*P* = 0.002).

**Fig 6.**
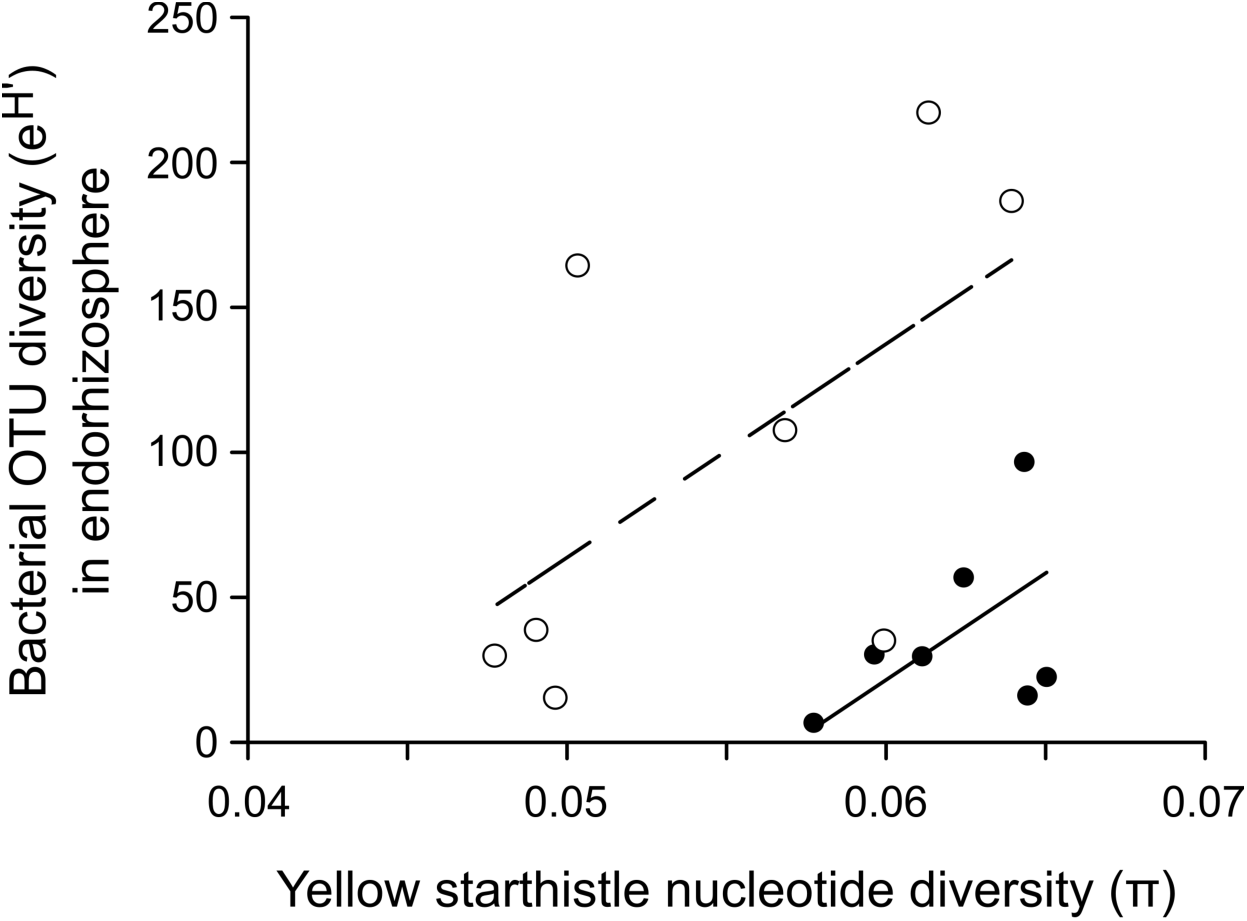
Bacterial diversity (e^H’^) in endorhizosphere samples pooled by sampling location, as a function of the genetic diversity among plants at the same sites (calculated as the average proportion of pairwise nucleotide differences between alleles (π) at variable positions in the genome; from Barker *et al.*, 2017). Lines show significant positive relationships (linear model: *P* = 0.02) between microbial and plant diversity at sampling locations in both the native range (open symbols, dashed line) and the invaded range (closed symbols, solid line).

## DISCUSSION

Introduced plants will encounter a variety of novel species interactions as they establish across biogeographic regions. For many plants, severe invasions are associated with more favorable interactions with soil microbial communities (Andonian & Hierro, 2011; Dawson & Schrama, 2016). We found that bacterial microbiomes of invading yellow starthistle were unique in composition and lower in diversity relative to bacterial microbiomes of native plants, differences that persisted within plant compartments and across variation in plant genetic diversity.

As observed in other species, bacterial communities differed most among plant compartments (Bulgarelli *et al.*, 2013; Vandenkoornhuyse *et al.*, 2015). The numbers and diversity of taxa within each compartment were similar in magnitude to those reported in other studies of prokaryotic 16S sequences, e.g. from Agavaceae (Coleman-Derr *et al.*, 2016), Brassicaceae (Bodenhausen *et al.*, 2013), Cactaceae (Fonseca-García *et al.*, 2016), and other Asteraceae (Leff *et al.*, 2016). The dominant phyla were Proteobacteria, Actinobacteria, Bacteroidetes, and Firmicutes, which are also characteristic of plant-associated bacterial communities surveyed to date (Bulgarelli *et al.*, 2013). The exception was the leaf endosphere, where a paucity of sequences relative to controls suggests that persistent chloroplast contamination obscured low frequency endophytes, despite our development of an Asteraceae-specific PNA (FitzPatrick *et al.*, submitted). A targeted survey is needed to better characterize this compartment (e.g. qPCR, (Fierer et al., 2005).

Notably, diversity was approximately twice as high in the endorhizosphere as in the ectorhizosphere. Current reviews have concluded that root endosphere communities are typically less diverse than those in the ectorhizosphere (Bulgarelli *et al.*, 2013; Vandenkoornhuyse *et al.*, 2015). Our root collections were washed but not surface sterilized and may represent some of the rhizoplane/rhizosphere in addition to the endosphere, elevating our estimates of diversity. It is also possible that yellow starthistle deviates from initially reported patterns, which have also been challenged by other recent studies (Fonseca-García *et al.*, 2016; Leff *et al.*, 2016).

Within compartments, community composition was consistently different between samples from native and invaded ranges and included shifts in taxa across all major groups. Our native range samples represented a larger geographic area and spanned distinct genetic subpopulations of yellow starthistle, but native range sites clustered together in overall community composition and there was little evidence of individual site differences within ranges. Between ranges, the diversity of OTUs was lower in the invaded range, a pattern that was dominated by lower evenness of OTUs in both the phyllosphere and ectorhizosphere, and by lower richness of OTUs in the endorhizosphere. Thus invading plants were more strongly dominated by a few taxa at high relative abundance on root and leaf surfaces, and harbored fewer bacterial taxa in their root endophytic communities.

We observed a significant positive association between root microbial diversity and genotypic diversity among plants at the population scale. This association was strongest in the endorhizosphere, the only endophytic compartment in our analysis, consistent with plant genotype having the largest influence on microbial taxa colonizing within the plant itself (Bulgarelli *et al.*, 2012; Lundberg *et al.*, 2012; Lebeis *et al.*, 2015). Within-species plant genotype effects have been observed previously and may interact with the effect of environment to shape microbial communities (Peiffer *et al.*, 2013; terHorst & Zee, 2016). Interactions between plant and microbial diversity could be particularly important for invasive species, where genetic bottlenecks during establishment and range expansion can reduce genetic diversity among plants (Dlugosch & Parker, 2008; Excoffier *et al.*, 2009; Uller & Leimu 2011). Nevertheless, it appears that genotype effects are often minor relative to site effects (Lundberg *et al.*, 2012; Peiffer *et al.*, 2013; Bodenhausen *et al.*, 2014; Bulgarelli *et al.*, 2015), and we found that genotypic effects were evident only within region and did not explain microbial diversity differences between regions.

The microbial differences that we observed across ranges are similar to those observed at continental scales (Bodenhausen *et al.*, 2013; Peiffer *et al.*, 2013). A variety of factors may explain these patterns, particularly abiotic differences (Fierer & Jackson, 2006; Bulgarelli *et al.*, 2013; Nemergut *et al.*, 2013; Vandenkoornhuyse *et al.*, 2015). Soil type appears to have a strong influence on microbial communities (e.g. Bulgarelli *et al.*, 2012; Lundberg *et al.*, 2012), and is known to differ broadly across yellow starthistle’s range (Hierro *et al.*, 2017). In addition, populations in California are at the warm and dry extreme of yellow starthistle’s climatic niche (Dlugosch *et al.*, 2015), and our sampling was conducted at the end of a period of severe drought (Griffin & Anchukaitis, 2014; Diffenbaugh *et al.*, 2015), which could have amplified microbial differences related to climate (Schrama & Bardgett, 2016). Interestingly, a recent study of grassland plants found that microbial diversity increased under drought, whereas we found reduced diversity in the drought-affected range (terHorst *et al.*, 2014).

Importantly, yellow starthistle’s invasion could also cause changes to microbial communities. Species invasionss have been shown to alter microbial composition over short timescales (Collins *et al.*, 2016; Gibbons *et al.*, 2017), though long term effects are less clear (Lankau, 2011; Day *et al.*, 2015). Yellow starthistle invasions are denser than populations in the native range by an order of magnitude or more (Uygur *et al.*, 2004; Andonian *et al.*, 2011), and include lower diversity of plant species overall (Seabloom *et al.*, 2003; Zavaleta & Hulvey, 2004; D’Antonio *et al.*, 2007). Low plant diversity can depress the diversity of microbes in the environment and in plants (Garbeva *et al.*, 2004; Schnitzer *et al.*, 2011; Coleman-Derr *et al.*, 2016). Invading plants in California also grow to larger size, and could allow particular strains to proliferate if size increases were gained at a cost to defenses (Huot *et al.*, 2014; Dlugosch *et al.*, 2015). These possibilities mean that invasions could be a cause rather than an effect of weaker plant-soil interactions for invasive species in their introduced ranges (Kulmatiski *et al.*, 2008; Dawson & Schrama, 2016), reinforcing a growing need for explicit observational and experimental tests that disentangle associations between microbial diversity and plant density, plant community diversity, and environmental gradients (Dawson & Schrama, 2016).

To date, few studies have compared microbial community compositionl between the native and introduced ranges of invasive plants. McGinn and colleagues (2016) reported no differences in the diversity of mutualistic fungal taxa associated with the roots of multiple species of European *Trifolium* introduced to New Zealand, despite more favorable soil interactions in the invasions (McGinn et al., 2018; but see Parker & Gilbert, 2007. Johansen and colleagues (2017) found *increased* diversity of fungal communities on the roots of European *Ammophila arenaria* invading Australia and New Zealand, though there appear to be no differences in interactions with soil microbial communities in its invasions (in North America; Beckstead & Parker, 2003). Gundale and colleagues explored the potential contribution of fungal endophyte communities to more favorable (negative) soil interactions observed in introduced plantations of lodgepole pine (*Pinus contorta*) from North America (Gundale *et al.*, 2014; 2016). For lodgepole pine, microbial communities differed among several global regions examined, but there was no consistent pattern of loss of potential fungal pathogens or gain of mutualists in the introductions, and it remains unclear what part of the soil community is responsible for observed differences in interactions across ranges (Gundale *et al.*, 2016). Reinhart and colleagues (2010) focused specifically on *Phythium* fungal pathogens and quantified their virulence on North American *Prunus serotina* introduced to Europe. They found that the most virulent strains occurred only in the native range, consistent with benefits to invading plants escaping components this specific pathogenic group. In the only microbiome comparison that included bacterial taxa, Finkel and colleagues (Finkel *et al.*, 2011; 2016) explored the phyllosphere community of multiple species of *Tamarix* in native and introduced parts of their ranges, finding that microbial communities were in general most strongly structured by geographic region.

Our study is the first to find consistent differences in the microbiomes of native and invading plants which coincide with fitness differences in plant interactions with soil communities. Among the few invader microbiome studies to date, ours is unusual in focusing on the bacterial community. Fungi have historically received more attention for their fitness effects on plants, but bacteria can also play a critical role both as pathogens and mutualists (Haney *et al.*, 2015; Vandenkoornhuyse *et al.*, 2015; Herrera Paredes & Lebeis, 2016). For yellow starthistle, previous experiments have demonstrated that fungal communities are not responsible for more favorable conditions in the invaded range (Hierro *et al.*, 2017), and our findings indicate that bacterial communities warrant further investigation as the potential source of these differences. The fitness effects of our specific OTUs are unknown, however, and identifying the bacterial OTUs that accumulate during interactions with plants would help to elucidate important pathogenic or mutualistic taxa, and allow field surveys to explicitly test hypotheses that these strains are lost or gained in the invaded environment.

We have previously argued that yellow starthistle has benefitted from the historical loss of plant competitors in California (Dlugosch *et al.*, 2015). Disturbance is critical for yellow starthistle establishment, and functionally similar native species compete well against it in experiments; however, key competitors have been lost from the ecosystem due toperturbations prior to yellow starthistle invasion (Zavaleta & Hulvey, 2004; Hooper & Dukes, 2010; Hierro *et al.*, 2011, 2017; Hulvey & Zavaleta, 2012). Any benefits of altered bacterial communities could be independent of competition with native plant species, but these factors might also interact. Increased density due to a lack of competition could have reduced plant-associated microbial diversity, as described above. Yellow starthistle experiences negative plant-soil feedbacks across generations (Andonian *et al.*, 2011), however, and the build up of high plant densities is therefore unlikely to generate more favorable soil interactions. Alternatively, the historical loss of native species diversity in California (D’Antonio *et al.*, 2007) could have resulted in the loss of associated microbial diversity (Garbeva *et al.*, 2004; Schnitzer *et al.*, 2011; Coleman-Derr *et al.*, 2016), generating particularly strong opportunities for invasion into a system with both reduced plant competition and reduced pathogen diversity. Microbial surveys of remnant native communities, as well as across densities of yellow starthistle would facilitate tests of alternative hypotheses for interacting effects of plant and microbial diversity, and it may be informative to explore microbial communities preserved on native plant specimens pre-dating the extensive invasion of yellow starthistle into this region.

## Conclusions

We found consistent differences between native and invading yellow starthistle plants in their bacterial microbiomes. These differences were robust to additional variation associated with plant compartment and the diversity of plant genotypes. Invaded range microbiomes differed in composition across major taxonomic groups, and harbored a lower diversity of bacteria, including reduced evenness on the surface of leaves and roots and reduced richness of root endophytes. We suggest that bacteria could be the source of more favorable microbial interactions that have been observed in this invasion. Our findings also raise questions about 1) whether lower bacterial diversity is a feature of the invaded environment or whether it is caused by the invasion itself, and 2) how differences in the microbial community might have interacted with other changes in plant competitive interactions and the abiotic environment to affect plant fitness. These questions highlight the need for additional studies that compare microbial communities (including bacteria) associated with native and invading populations, that couple microbial community identification with plant-soil feedback and fitness experiments, and that examine the interaction of environment, plant diversity, and plant density on microbial communities and their fitness effects on plants.

## ACKNOWLEDGMENTS

We thank D. Lundberg and S. Lebeis for helpful discussions regarding sampling design; G. Reardon and C. Chandeyson for assistance with field collections; E. Arnold, J. Aspinwall, J. Braasch, E. Carlson, K. Hockett, P. Humphries, T. O’Connor, M. Schneider, J. U’Ren, and N. Zimmerman for 16S library preparation assistance and discussion; A. Gerritsen, D. New and staff at iBEST for assistance with sequencing; K. Andonian, J. Hierro, and 3 reviewers for helpful feedback on the manuscript. We are particularly indebted to C.E. Morris for hosting the European sample processing at INRA Station de Pathologie Végétale, Montfavet, France. Data collection and analyses performed by the IBEST Genomics Resources Core at the University of Idaho were supported in part by NIH COBRE grant P30GM103324. This study was supported by USDA grant 2015-67013-23000 to K.M.D., D.A.B., and S.M.S.

## AUTHOR CONTRIBUTIONS

P.L-I., D.A.B., and K.M.D. designed the study. P.L-I. and J.H. collected the samples with assistance from H.S., S.M.S., and S.R.W. P.L-I conducted the microbial sequencing and bioinformatics. P.L-I, S.R.W., and K.M.D. analyzed the data. P.L-I and K.M.D. wrote the manuscript, which was edited by all authors.

## SUPPORTING INFORMATION

**Fig. S1.**
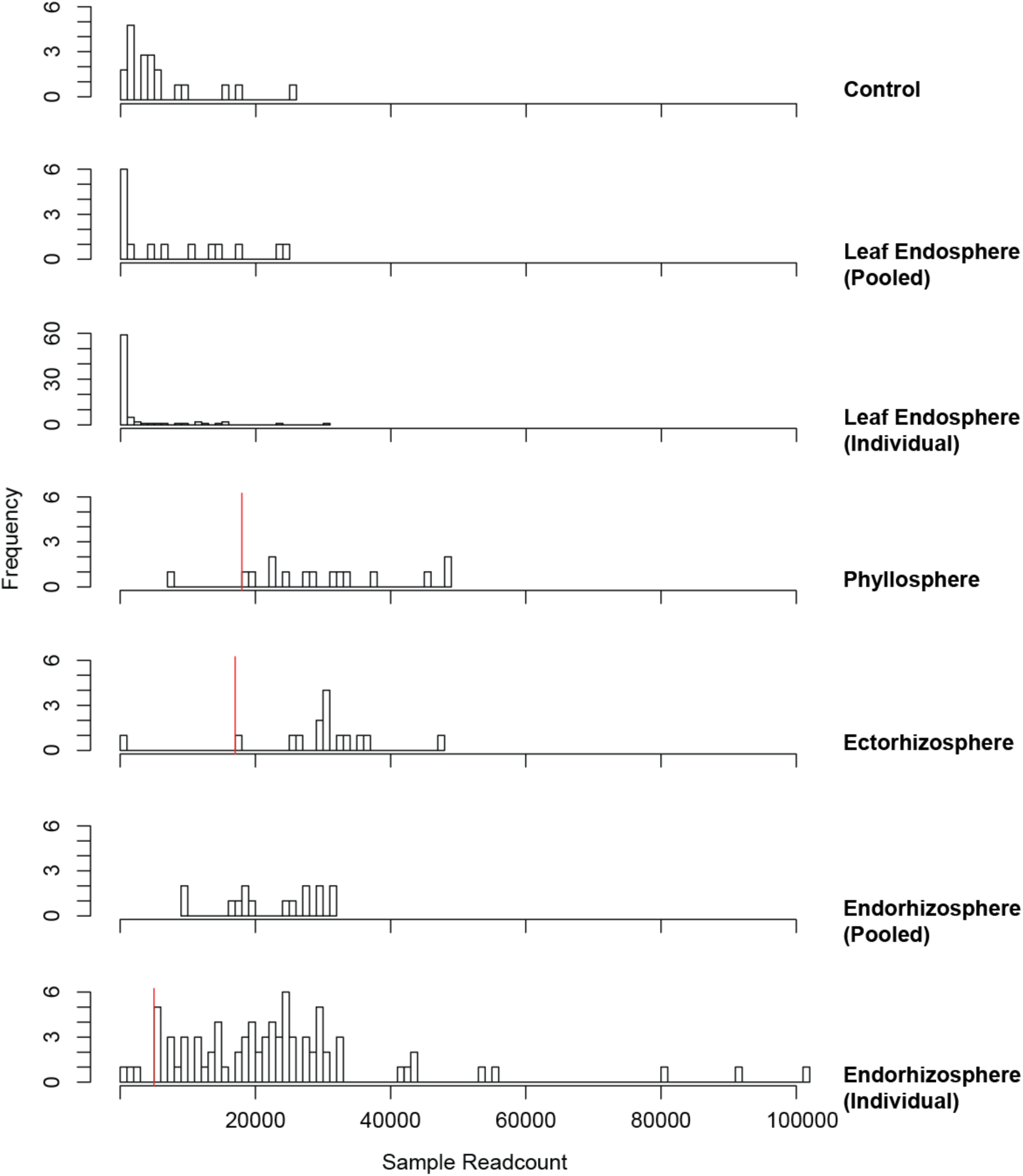
Frequency distributions of un-rarefied read counts for samples from all four plant compartments (native and invading population samples combined), as well as control (blank) samples. Vertical red lines indicate thresholds for rarefaction where relevant.

**Fig. S2.**
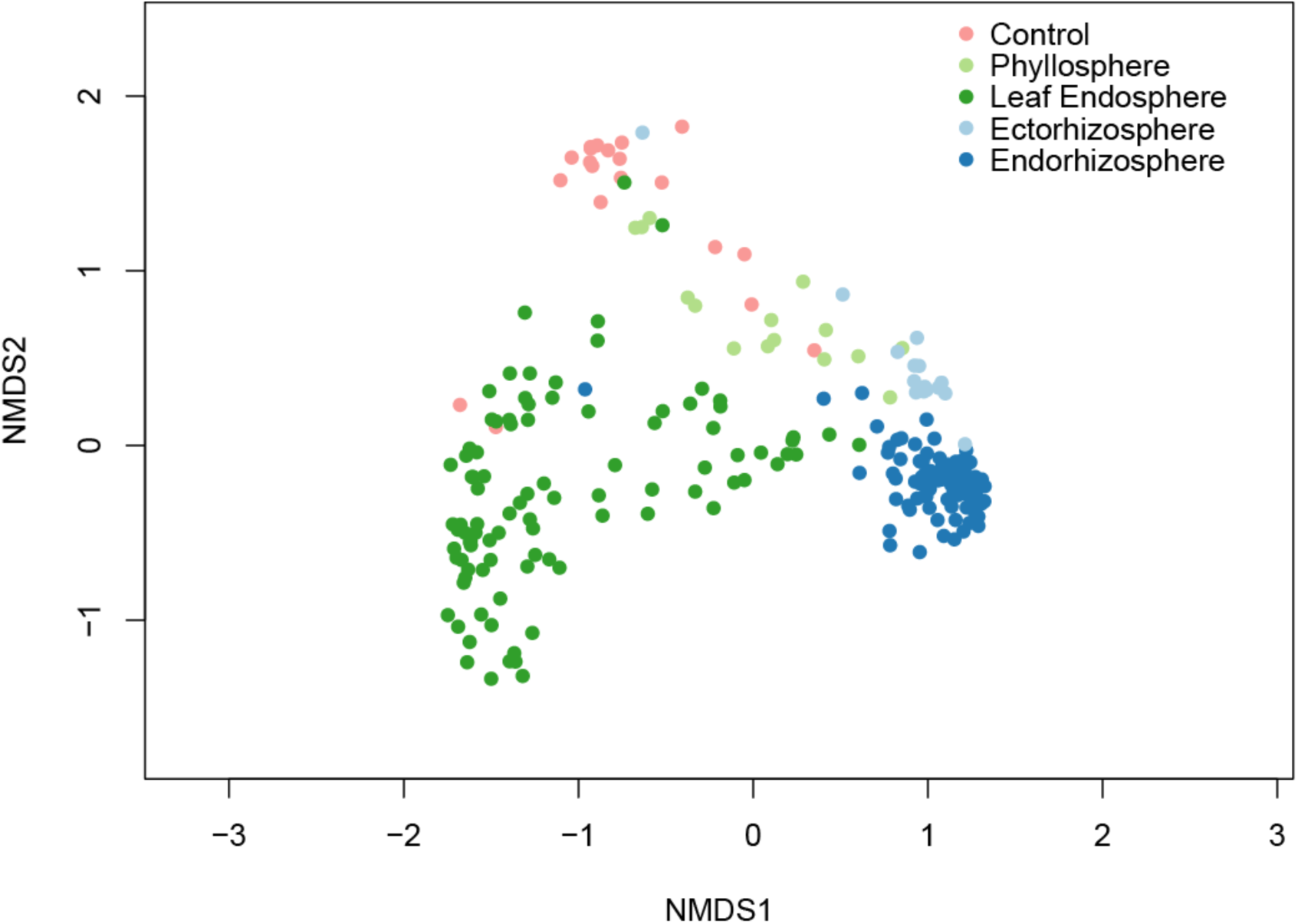
NMDS plot of bacterial OTU composition in unrarefied control, phyllosphere, leaf endosphere, ectorhizosphere, and endorhizosphere samples (stress = 0.10).

**Fig. S3.**
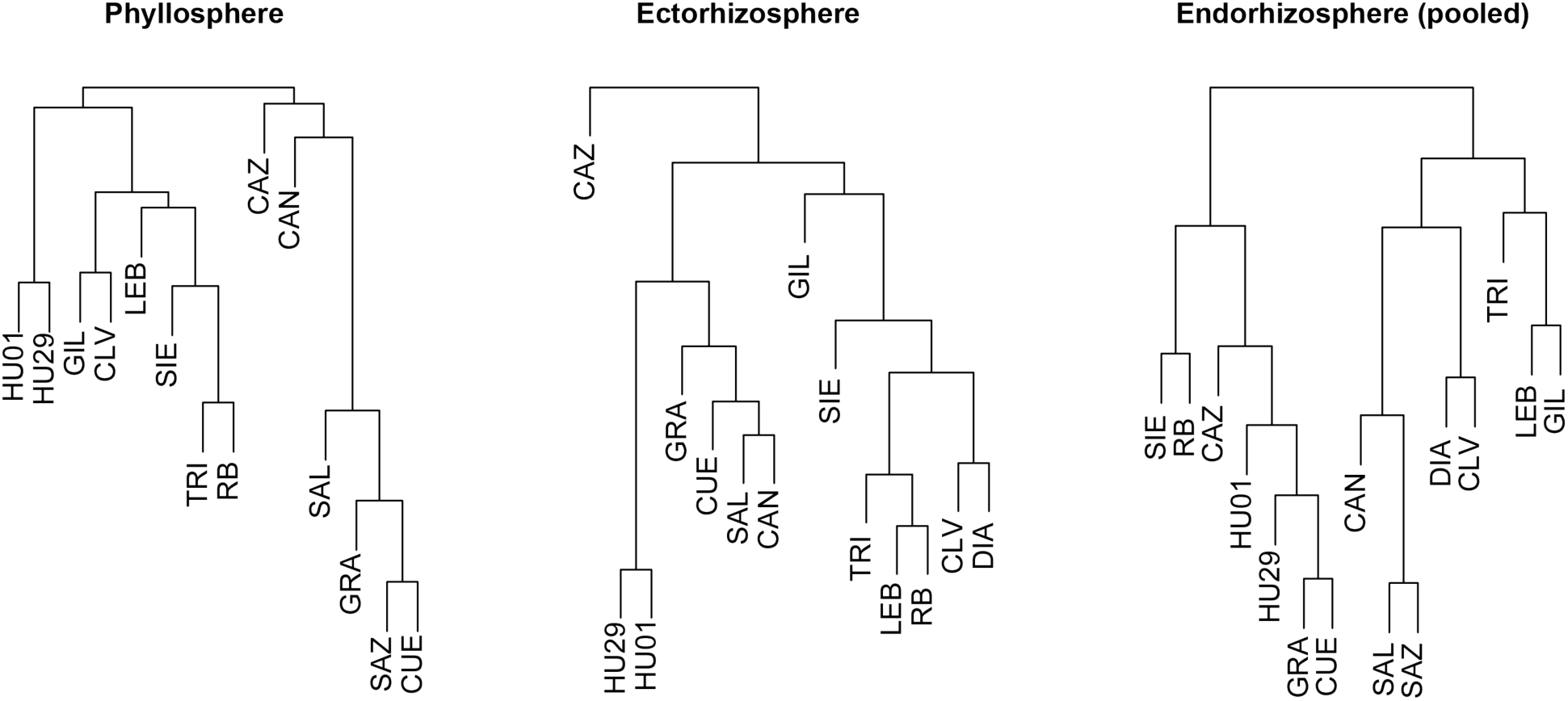
Yellow starthistle populations clustered according to Bray-Curtis dissimilarity between rarefied, square-root transformed pooled samples from three plant compartments.

**Fig. S4.**
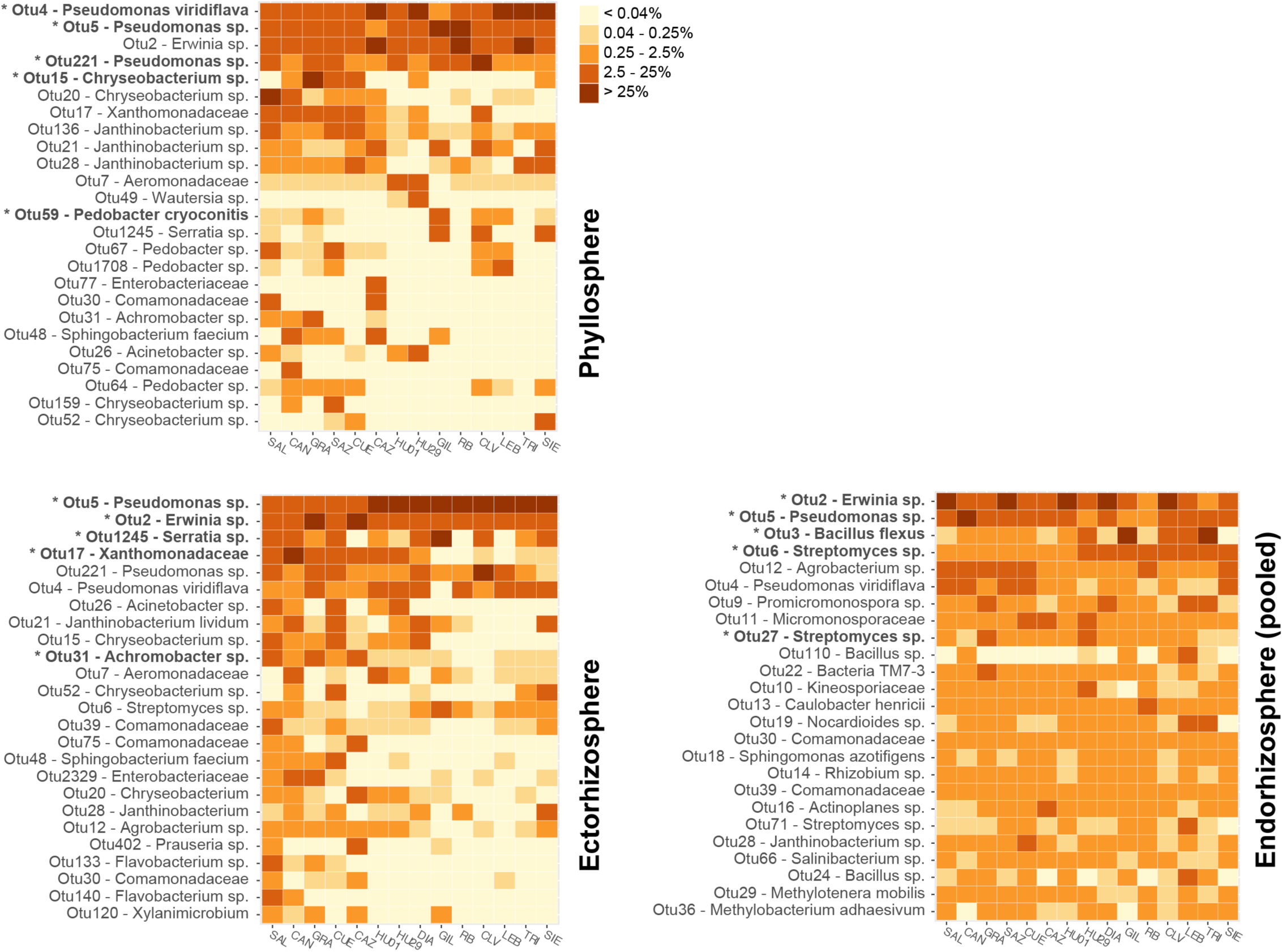
Heat maps showing the fraction of reads mapping to each of the top 25 OTUs in rarefied bulk samples from three plant compartments. The top five OTUs contributing to the first principal component separating native and invaded samples in each compartment are indicated with an asterisk. Taxonomic assignments for each OTU are given to the lowest taxonomic rank identified.

**Fig. S5.**
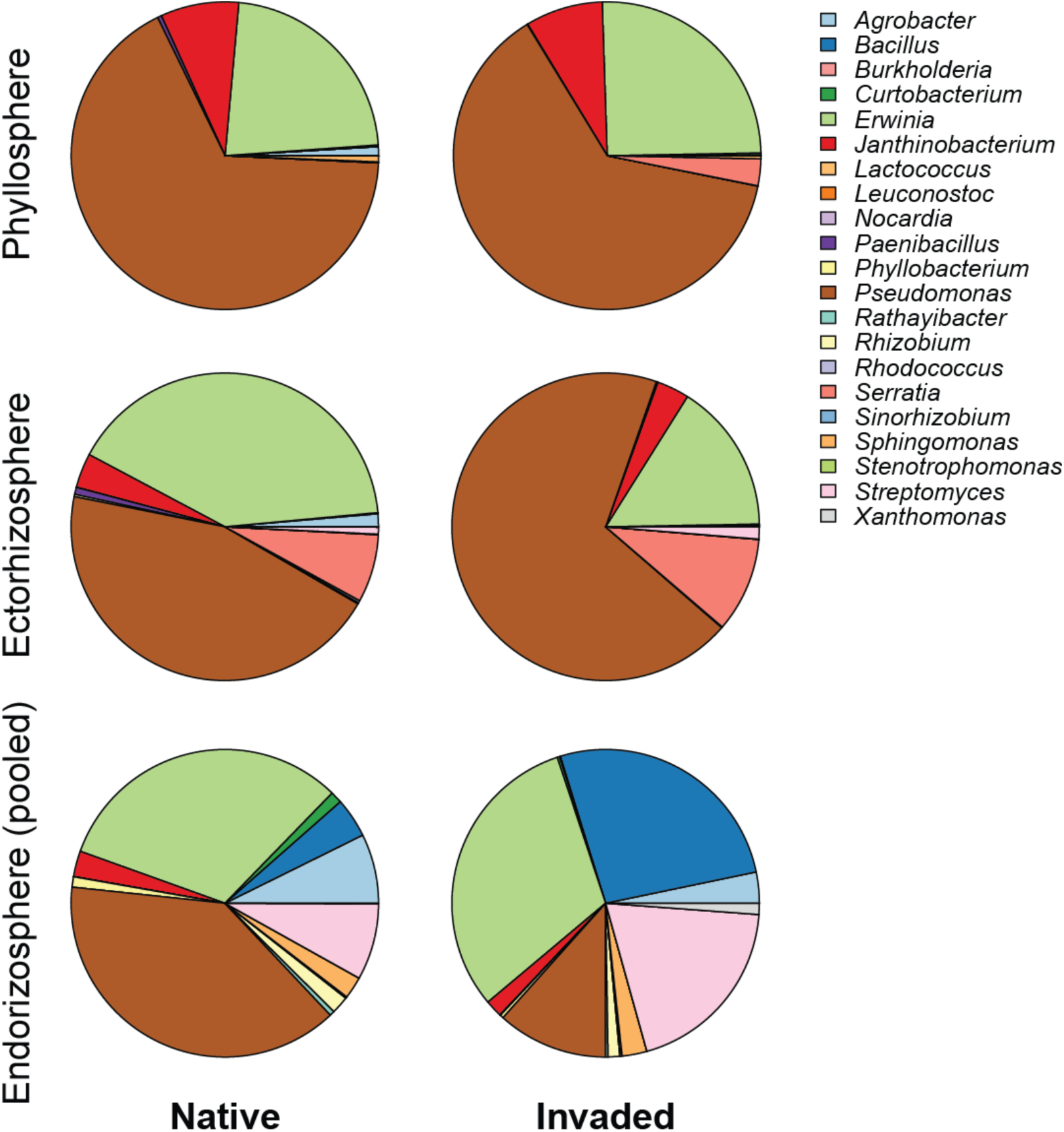
Proportion of rarefied read counts assigned to pathogen-containing genera in all samples according to plant compartment and range.

**Table S1.** Collecting information for populations sampled in this study, including designation codes, localities, coordinates, sampling dates, and specimen accession numbers of plant collections at ARIZ.

| Code | Lat. | Long. | Locality | Date | ARIZ specimen numbers |
| --- | --- | --- | --- | --- | --- |
| SAL | N40.99112 | W005.65831 | Avenida de la Merced, ~1.5km N of Salamanca. Dirt path between road and field, on E side of road. | 22-Jun-15 | 426137, 426032, 425636, 426125, 426034, 426033, 426035, 426126, 426140, 426068, 426127, 426128, 426129, 426130, 426131, 426132, 426133, 426134, 426135, 426136, 426138, 425635, 426139 |
| CAN | N41.00085 | W004.89715 | Junction AV800 & road to Canales. On E side of Canales road, in the narrow strip between road and field edge. | 22-Jun-15 | 425643, 426065, 426064, 426063, 426055, 426054, 425646, 426067, 426053, 426052, 425642, 426066, 426051, 425641, 426062, 426061, 426059, 426058, 426060, 426057, 426056 |
| GRA | N37.26843 | W003.66488 | GR300 ~100m from junction w 213, ~20m up dirt path to the E side of the road. Weedy waste area near edge of olive orchard. | 23-Jun-15 | 425716, 425715, 425714, 425713, 425739, 425738, 425737, 425736, 425735, 425734, 425733, 425732, 425728, 425740, 425729, 425711, 425702, 425703, 425704, 425705, 425706, 425707, 425708, 425709, 425710 |
| SAZ | N39.83529 | W002.50997 | Gravel road by W side of CM3118 (road to Villares del Saz), just next to underpass under A3. Dirt farming path (for tractors) on S side of road, between sunflower fields. | 24-Jun-15 | 425678, 425741, 425679, 425731, 425680, 425681, 425682, 425683, 425684, 425685, 425757, 425755, 425753, 425751, 425749, 425746, 425744, 425743, 425758, 425756, 425752, 425750, 425748, 425747, 425745, 425742 |
| CUE | N40.12939 | W002.13880 | CM2105 between Cuenca & Tragacete ~8km W of Cuenca. Gravel road to the N of 2105, between field and river. | 24-Jun-15 | 425424, 425421, 425420, 425423, 425425, 425426 |
| CAZ | N43.74958 | E003.77119 | D113, ~1km from junction w 986, near Cazevielle. Flat, gravelly turnout & intersection w unpaved road, & gravelly roadsides. | 20-Jun-15 | 425691, 425690, 425971, 425688, 425689, 425972, 425692, 425970, 425693 |
| HU01 | N47.18240 | E18.08928 | Unnamed dirt roads about 200m NW of E66/8, running parallel. Access by perpendicular unnamed paved road. | 7-Jul-15 | 425655, 425660, 425661, 425653, 425652, 425658, 425657, 425656, 425654, 425631, 425687, 425634, 425633, 425686, 425630, 425628, 425629, 425627, 425662 |
| HU29 | N47.31785 | E21.03385 | Margin of irrigation ditch bordered by hay and sunflower fields on one side and service road parallel to main road (E60/4) on the other side. Site to the south of E60/4. | 8-Jul-15 | 425730, 425677, 425676, 425640, 425675, 425674, 425673, 425726, 425725, 425724, 425723, 425722, 425721, 425720, 425719, 425718, 425717, 425727 |
| DIA | N37.86275 | W121.98003 | Hillside to the West of parking lot at the end of Green Valley Road. | 8-Jun-15 | 425394, 425400, 425401, 425402, 425403, 425404, 425405, 425406, 425407, 425408, 425409, 425115, 425381, 425410, 425382, 425383, 425384, 425385, 425386, 425387, 425388, 425389, 425390, 425391, 425392 |
| GIL | N37.03389 | W121.53611 | Disturbed grassy area adjacent to turn out on South side of Leavesley Rd. near intersection with New Ave. Grassy patch between road and tilled field. | 7-Jun-15 | 425374, 425373, 425380, 425379, 425378, 425377, 425376, 425375, 425124, 425123, 425122, 425121, 425119, 425120, 425118, 425114, 425113, 425399, 425112, 425395, 425111, 425398, 425397, 425396 |
| RB | N40.27085 | W122.27103 | Lightly grassy patch to north-west of rest stop (Herbert S. Miles rest area) between Cottonwood and Red Bluff on southbound I-5. | 9-Jun-15 | 425372, 425371, 425370, 425369, 425368, 425418, 425365, 425415, 425367, 425366, 425416, 425417, 425419 |
| CLV | N36.91603 | W119.79341 | Gully on eastern side of the Yosemite fwy (41) adjacent to olive orchards. | 6-Jun-15 | 426046, 426045, 426044, 426043, 426042, 426036, 426037, 425637, 426038, 426039, 426040, 426041, 425639, 426026, 426027, 426028, 426029, 426030, 426031, 425638 |
| LEB | N34.82844 | W118.87369 | Cal trans maintained, weedy area adjacent to SW of rest stop off of south bound I-5. | 5-Jun-15 | 425968, 425695, 425969, 425966, 425967, 426047, 425694, 426050, 426049, 426048 |
| TRI | N37.46159 | W119.79399 | Strip of grassland along south side of Hwy 49 at intersection with Triangle road. Strip between road and fenced off rangeland. | 11-Jun-15 | 425116, 425117, 425414, 425412, 425413, 425411, 425393 |
| SIE | N38.78162 | W120.41639 | Site upslope of Weber Mill rd. about ¼ mile from the turnoff from Ice House road. | 10-Jun-15 | 425648, 425647, 425649, 425644, 425645 |

**Table S2.**
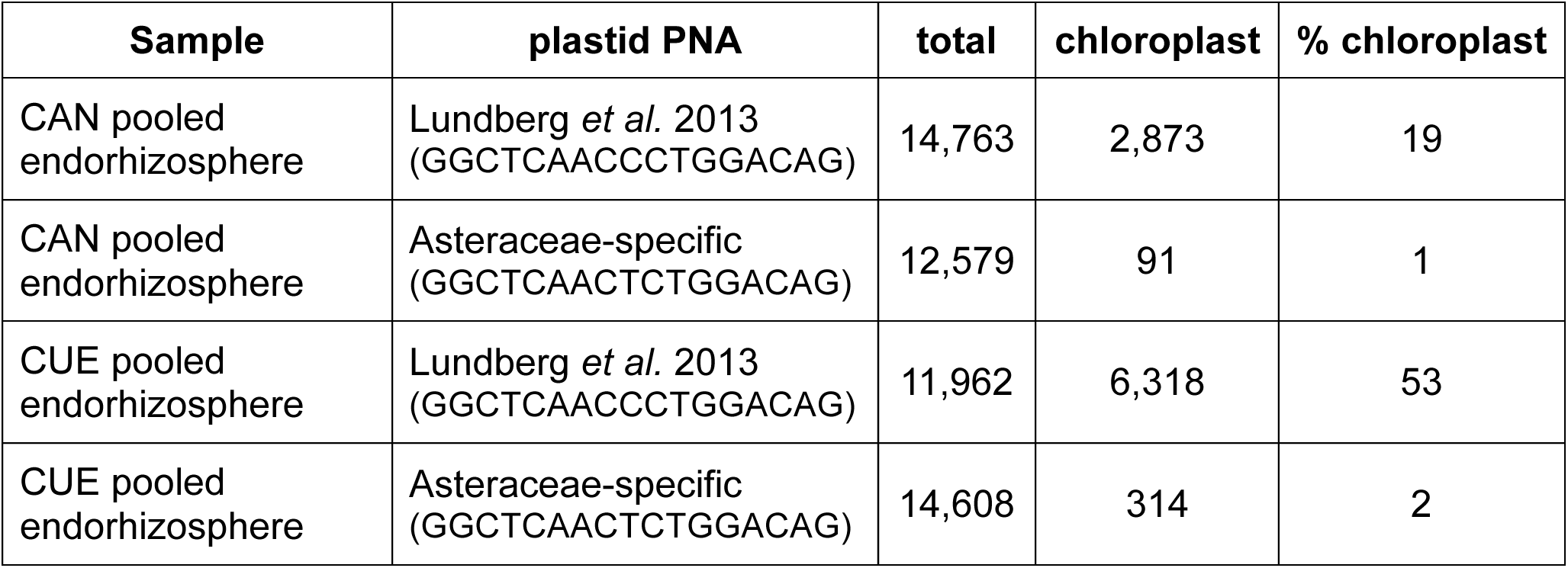
Comparison of host chloroplast blocking levels achieved by different PNA clamps in pooled endorhizosphere samples from two populations. Total read counts, number of reads mapping to host chloroplast, and host chloroplast reads as percentage of total reads are reported for each sample.

**Table S3.** Loading by top 10 individual OTUs along first principal component axis from analysis of Hellinger transformed data matrices for three plant compartments. Column A: individual OTU unique identification number; column B: individual OTU lowest taxonomic assignment; column C: axis loading per individual OTU.

| Phyllosphere |  |  |
| --- | --- | --- |
| A | B | C |
| 15 | *Chryseobacterium* | 0.51 |
| 4 | *Pseudomonas viridiflava* | 0.41 |
| 17 | Xanthomonadaceae | 0.32 |
| 221 | *Pseudomonas* | 0.27 |
| 2 | *Erwinia* | 0.26 |
| 20 | *Chryseobacterium* | 0.22 |
| 136 | *Janthinobacterium* | 0.20 |
| 7 | Aeromonadaceae | 0.18 |
| 49 | *Wautersiella* | 0.16 |
| 31 | *Achromobacter* | 0.16 |

| Ectorrhizosphere |  |  |
| --- | --- | --- |
| A | B | C |
| 5 | *Pseudomonas* | 0.62 |
| 17 | Xanthomonadaceae | 0.44 |
| 31 | *Achromobacter* | 0.24 |
| 26 | *Acinetobacter* | 0.24 |
| 15 | *Chryseobacterium* | 0.20 |
| 48 | *Sphingobacterium faecium* | 0.15 |
| 75 | Comamonadaceae | 0.14 |
| 4 | *Pseudomonas viridiflava* | 0.13 |
| 221 | *Pseudomonas* | 0.13 |
| 2329 | Enterobacteriaceae | 0.12 |

| Endorhizosphere |  |  |
| --- | --- | --- |
| A | B | C |
| 2 | *Erwinia* | 0.56 |
| 3 | *Bacillus flexus* | 0.41 |
| 5 | *Pseudomonas* | 0.41 |
| 6 | *Streptomyces* | 0.17 |
| 10 | Kineosporiaceae | 0.15 |
| 11 | Micromonosporaceae | 0.13 |
| 4 | *Pseudomonas viridiflava* | 0.13 |
| 27 | *Streptomyces* | 0.11 |
| 13 | *Caulobacter henricii* | 0.10 |
| 23 | Actinomycetales | 0.10 |

